# Shipwrecks can mirror predator assemblages of pelagic pinnacles but may lack the trophic balance of natural reef environments

**DOI:** 10.1101/2025.09.10.675479

**Authors:** Scarlett R. Taylor, Gavin Miller, Piers Baillie

**Affiliations:** Global Reef, 45/1 M3 Koh Tao, Surat Thani, Thailand, 84360

**Keywords:** artificial reefs, fish assemblages, trophic structure, Bayesian inference, marine conservation

## Abstract

Artificial reefs are widely deployed to enhance fish populations, yet their capacity to replicate the ecological roles of natural reef systems remains uncertain, and comparisons are often limited to nearby fringing reefs. Such assessments may overlook how structural features influence ecological outcomes. This study compared fish assemblages across shipwrecks, pelagic pinnacles, and fringing reefs in the Gulf of Thailand to evaluate how artificial structures align with natural patterns of community structure and trophic composition. Natural reef sites were selected to represent contrasting structural profiles, with pinnacles characterized by high vertical relief and fringing reefs by lower-relief slopes. This design enabled assessment of whether high-relief artificial structures, such as shipwrecks, more closely reflect fish communities found on similarly structured natural reefs. Bayesian multivariate models applied to 350 underwater surveys from 14 sites showed that shipwrecks resembled pelagic pinnacles in supporting mesopredators and higher trophic level predators, but had consistently lower representation of grazers and invertivores relative to both natural reef types. These results indicate functional divergence likely driven by reduced benthic complexity and limited forage availability. While shipwrecks may support fisheries species and predator biomass, they do not fully function as ecological analogues of natural reefs. Findings highlight the importance of aligning artificial reef design with defined ecological objectives and considering structural context in management strategies across tropical reef systems worldwide.

## 1. INTRODUCTION

Artificial reef performance is typically evaluated by comparison with nearby fringing reef systems, yet these assessments may overlook how structural features influence ecological outcomes. Fringing reefs, which dominate nearshore tropical coastlines, are generally low in vertical relief and often embedded in sediment-rich, human-impacted environments (Bell & Galzin 1984, Letessier et al. 2019). In contrast, many artificial reefs, particularly shipwrecks, share greater physical similarity with offshore pelagic pinnacles, including steep vertical profiles, current exposure, and spatial isolation (Letessier et al. 2019, Paxton et al. 2020b, Galbraith et al. 2022). These structural features influence foraging opportunities, predator access, and planktonic input, potentially shaping fish community composition and trophic balance (White et al. 2007, Galbraith et al. 2023). Despite these parallels, few studies have examined whether shipwrecks more closely resemble the ecological characteristics of pinnacles, align instead with fringing reef communities, or represent a functionally novel assemblage.

Artificial reefs are deployed worldwide to mitigate habitat loss and enhance fish populations (Rilov & Benayahu 2000, Paxton et al. 2020a, Harvey et al. 2021, Baillie et al. 2024). Among the various designs, shipwrecks are frequently used due to their complex structure, tourism appeal, and availability through both intentional deployment and maritime accidents (Mattos & Yeemin 2020, Medeiros et al. 2022, Hickman et al. 2024). In Thailand, shipwreck deployment has been widespread since the launch of the national Artificial Reef Program in 1978, aimed at reducing diver pressure on natural reefs and supporting fisheries enhancement (Yeemin et al. 2006, Mattos & Yeemin 2020, Harvey et al. 2021, Baillie et al. 2024). The Gulf of Thailand contains three primary reef habitat types relevant to this study: nearshore fringing reefs that slope gradually toward sandy bottoms (Yeemin et al. 2009), deeper offshore pinnacles with substantial vertical relief, and a growing number of artificial structures such as shipwrecks (Mattos et al. 2023). Despite the prevalence of these artificial reefs, relatively few studies have assessed their ecological functioning, particularly their capacity to support balanced trophic assemblages and replicate the ecological roles of natural reef systems (Fowler & Booth 2012, Simon et al. 2013, Streich et al. 2017, Medeiros et al. 2022, Paxton et al. 2023, Schram et al. 2024).

Where evaluations do exist, artificial reefs, particularly shipwrecks, often support high total fish abundance and species richness compared to nearby natural reefs (Fowler & Booth 2012, Folpp et al. 2013, Schulze et al. 2020). However, these metrics can obscure differences in community composition and ecological function (Ross et al. 2016, Paxton et al. 2020a, Paxton et al. 2020b, Paxton et al. 2023). Predator biomass is frequently elevated, while lower trophic guilds such as grazers and invertivores tend to be underrepresented on artificial reefs (Fowler & Booth 2012, Schram et al. 2024). This imbalance may limit processes such as algal control, nutrient cycling, and trophic transfer, indicating that high fish abundance alone does not equate to ecological equivalence with natural reefs.

Such divergences are influenced by multiple ecological and structural factors, including vertical relief, habitat complexity, reef age, and proximity to natural systems (Rilov & Benayahu 2000, Simon et al. 2013, Streich et al. 2017, Paxton et al. 2023). Shipwrecks tend to be high-relief and spatially isolated, favouring aggregation of upper trophic level species while offering limited support for lower trophic guilds (Harvey et al. 2021, Medeiros et al. 2022, Schram et al. 2024). Over time, these dynamics may produce assemblages that diverge functionally and compositionally from adjacent natural reefs (Paxton et al. 2020b, Paxton et al. 2023, Sibley et al. 2023). Distinguishing superficial species-level similarity from broader ecological performance is therefore essential when evaluating artificial reef outcomes (Simon et al. 2013, Medeiros et al. 2022, Schram et al. 2024). Variation among natural reef types provides important context for interpreting artificial reef performance. Isolated pelagic pinnacles, often exposed to strong hydrodynamic flow, tend to support high densities of mesopredators and larger predators such as *Caranx*, *Lethrinus*, *Lutjanus*, and *Epinephelus* spp. (Letessier et al. 2019, Galbraith et al. 2022, Cresswell et al. 2023, Galbraith et al. 2023a, Galbraith et al. 2023b, Mattos et al. 2023). These physical features enhance planktonic input and nutrient delivery, increasing productivity at lower trophic levels and facilitating biomass accumulation at higher trophic levels (White et al. 2007, Galbraith et al. 2023). Pinnacles also exhibit substantial vertical relief and structural complexity, which provide refuge and foraging opportunities across multiple trophic guilds (Galbraith et al. 2023, Letessier et al. 2019).

In contrast, fringing reefs and other emergent coastal systems are shaped more strongly by local variation in benthic composition, hydrodynamic exposure, and human disturbance (Bell & Galzin, 1984, Letessier et al. 2019, Galbraith 2021). These environments tend to support more trophically balanced assemblages, with a greater relative abundance of grazers and invertivores that play key roles in reef resilience and benthic regulation (Koeck et al. 2014, Boaden & Kingsford 2015, Streit et al. 2015, Schram et al. 2024). Comparisons between artificial reefs and adjacent natural reefs, including fringing systems, have consistently shown that shipwrecks support higher densities of mid- to higher-trophic fish, such as *Lutjanus*, *Sphyraena*, *Plectorhinchus*, and *Epinephelus* spp., but reduced densities of lower trophic guilds (Fowler & Booth 2012, Paxton et al. 2020a, Medeiros et al. 2022, Baillie et al. 2024, Schram et al. 2024). These patterns likely reflect limited food availability for grazers and invertivores on artificial substrates, as well as differences in benthic microhabitats and structural complexity (Rilov & Benayahu, 2000, Schram et al. 2024).

This study builds on previous assessments of fish assemblages in the Gulf of Thailand (Mattos & Yeemin 2020, Harvey et al. 2021, Mattos et al. 2023, Baillie et al. 2024), with the aim of evaluating whether shipwrecks replicate the community structure and trophic composition of natural reef analogues or represent a functionally novel assemblage type. Unlike most prior studies, which have compared artificial reefs only to nearby fringing reefs, this work explicitly tests shipwreck similarity against two contrasting natural reef types: fringing reefs and pelagic pinnacles, to determine which they more closely resemble. Given their structural similarity to pelagic pinnacles, shipwrecks are hypothesised to support predator-dominated assemblages more closely aligned with pinnacle reefs than with fringing systems. However, artificial reefs may lack the food availability, benthic complexity, or microhabitats required to sustain lower trophic guilds such as herbivores and invertivores (Fowler & Booth 2012, Simon et al. 2013, Schram et al. 2024). These divergences may reflect differences in habitat structure, trophic filtering, or broader ecological constraints. Characterising such patterns is critical to evaluating the ecological role of artificial reefs in tropical marine systems.

By explicitly comparing fish assemblages across shipwrecks, pinnacles, and fringing reefs, this study evaluates whether shipwrecks more closely resemble a particular type of natural reef or support distinct assemblages. Patterns in functional group composition, including the relative abundance of grazers, invertivores, mesopredators, and higher trophic level predators, are used alongside species-specific responses to assess ecological similarity and divergence. Key design features such as vertical relief, surface complexity, and orientation are considered in interpreting assemblage patterns, given their known influence on artificial reef performance (Rilov & Benayahu 2000, Paxton et al. 2020a). These findings contribute to a broader understanding of how artificial reefs function across structurally varied reef contexts and are intended to inform ecologically grounded deployment strategies not only in Thailand, where shipwreck use continues to expand, but in other tropical regions pursuing artificial reef development to support reef fisheries and conservation goals (Paxton et al. 2020a, Baillie et al. 2024).

## 2. MATERIALS AND METHODS

### 2.1. Study Sites

Surveys were conducted across three reef categories: fringing reefs (n = 5 sites), pelagic pinnacles (n = 6 sites), and artificial reefs (shipwrecks; n = 3 sites), totalling 14 individual sites (Figure 1). All sites were located around Koh Tao (10.10980° N, 99.81180° E) in the Gulf of Thailand, the largest semi-enclosed area in the central Indo-Pacific (Mattos et al. 2023). Koh Tao lies approximately 65 km offshore and provides access to a range of reef structures for comparative study. The three shipwrecks studied were decommissioned Royal Thai Navy vessels: the landing craft *HTMS Sattakut*, which was deployed in June 2011, and the two fast-attack vessels *HTMS Supharin* and *HTMS Hanhak Sattru*, both deployed in September 2023. Each site was surveyed repeatedly over the study period, yielding a total of 350 fish surveys: 150 on fringing reefs, 130 on pinnacles, and 70 on shipwrecks. Repeated surveys were performed at predetermined locations within each reef type to ensure consistency across dives.

**Figure 1.**
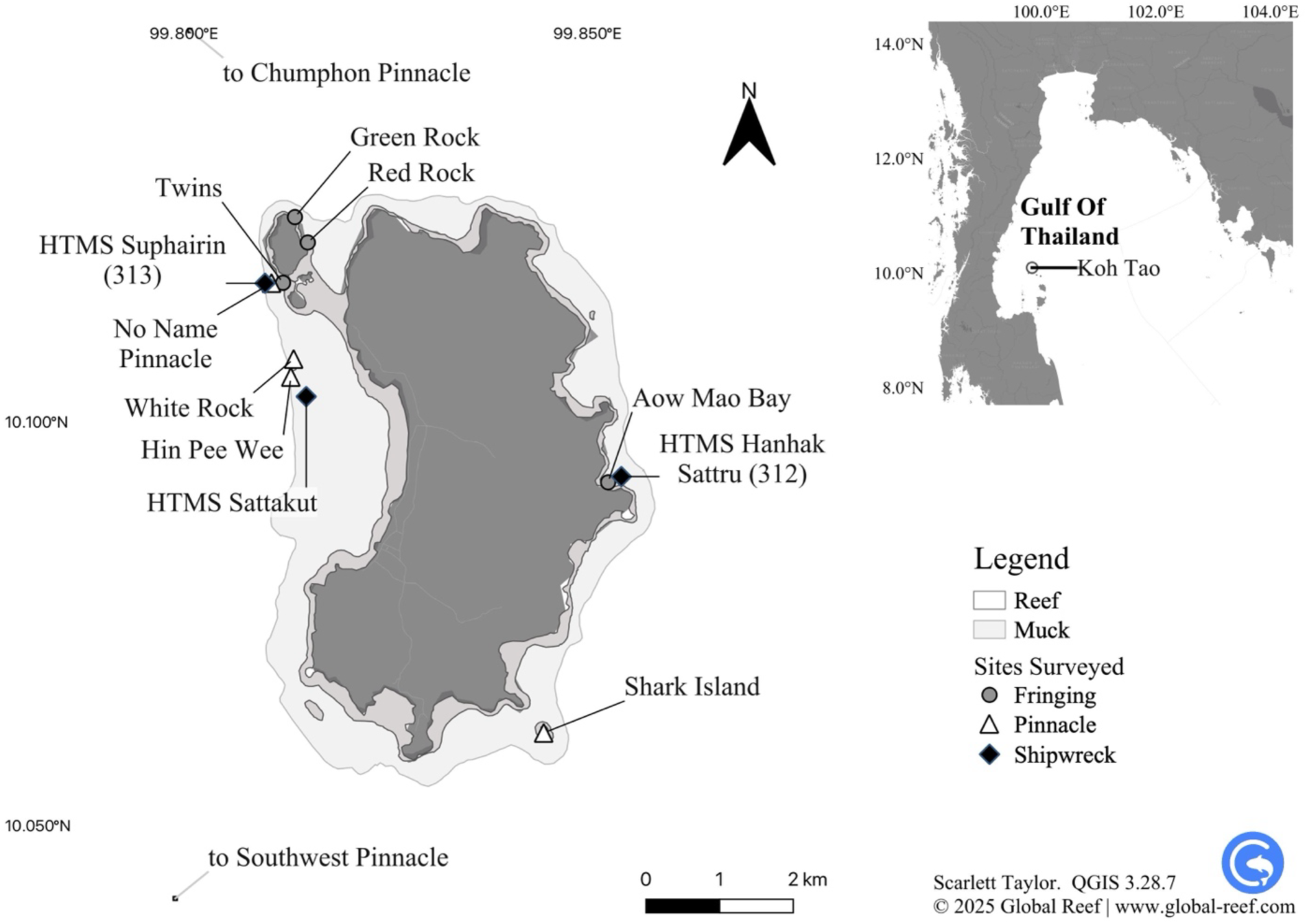
Dive sites surveyed around Koh Tao in the Gulf of Thailand (10.10980°N, 99.81180°E).

### 2.2. Survey Protocol

All surveys followed standardised underwater census (UVC) methodology. Divers conducted eight minute timed swim surveys using a predetermined survey species list (Table S1), which encompasses multiple functional groups (grazers, invertivores, mesopredators and higher trophic level predators) and ensures consistency in species identification and data collection. Eight minutes was selected as the survey duration based on multiple pilot surveys conducted on the *HTMS Sattakut,* where one full circuit of the wreck could be completed without overlap, minimizing double counting.

Each survey commenced from the same designated starting point on every dive site. The survey team consisted of two divers working in a buddy team on SCUBA. Divers maintained a position of 1-2 m away from each other at all times, and maintained a consistent swimming pace while ensuring they stayed level with their buddy. The observation field included all fish present within a 10-metre radius in the 180-degree field of view ahead, covering the left, forward, and right directions, and extending vertically to form a 180-degree three-dimensional arc above and below the divers. Divers swam slowly to maximise observation time and reduce disturbance to the fish. The estimated area covered during each survey was calculated by measuring the distance covered during multiple pilot timed-swim surveys on the HTMS *Sattakut*. All observed fish on the species list within the designated radius were recorded on waterproof slates. Data were recorded in real-time during the survey. Each new survey was recorded in a separate section of the slate to maintain data integrity.

### 2.3. Data Analysis

Data were analysed using Bayesian multivariate regression models, fit in R version 4.5.0 (2025-04-11) using the *brms* package (Bürkner 2017). Bayesian models were estimated using Markov chain Monte Carlo (MCMC) sampling implemented in Stan via *brms*. Data manipulation was performed using the *tidyverse*, and visualisations were generated with *ggplot2* (Wickham et al. 2019). The species were first grouped into functional groups – grazers, invertivores, mesopredators, and higher trophic level predators (hereafter HTLPs) – based on trophic ecology (Table S1) for analysis of broader ecological patterns and to facilitate interpretation of community-level responses. Separate multivariate models were fitted to evaluate differences in functional group abundance across reef classifications and, independently, across geographic zones. Subsequent species-specific models were used to examine finer-scale patterns within each functional group.

#### 2.3.1. Functional group analysis

To examine differences in the abundance of these functional groups across reef classifications, a Bayesian multivariate regression model was used. The model specified the joint response as: *mvbind*(*Grazer, Invertivore, Mesopredator, HTLP*) ∼ *Classification* + (1 + *Site*) allowing inference across groups while accounting for potential correlations in abundance patterns. Classification refers to the site type, either shipwreck, fringing reef or pelagic pinnacle. Shipwrecks were set as the reference group for reef classification.

Prior to model fitting, exploratory analyses were conducted to examine survey effort, distribution of counts, and the composition of functional groups across sites. Zero-inflation rates were calculated for each species, and species with infrequent but extreme counts (e.g. *Sphyraena* spp., *Gymnothorax* spp., *Tetraodon* and *Diodon* spp., all rays) were excluded due to their disproportionate influence on group-level patterns (<2% of observations removed). Functional group stability was assessed using coefficients of variation, and proportional group composition was assessed by site to evaluate consistency across reef types. These diagnostics informed data filtering and supported the robustness of the functional group classification. Zero-inflation was not evident in the remaining functional group data, while overdispersion was moderate, supporting the use of a negative binomial distribution.

Several candidate models were evaluated, including those with and without zero-inflation terms and with or without site-level random intercepts to account for repeated surveys. Model performance was compared using leave-one-out cross-validation (LOO-CV; Vehtari et al. 2017). The best-supported model included a negative binomial likelihood with random intercepts by site, capturing both overdispersion and hierarchical structure in the data. The response for each group *g*, at site *i*, survey *j*, was modelled as:

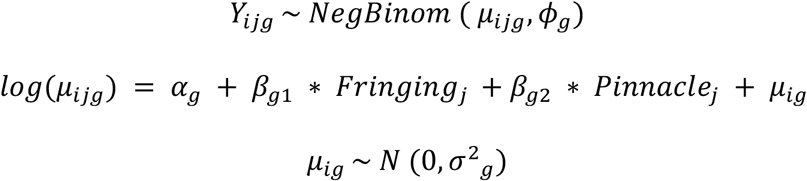

Here, *Y_ijg_* represents the count of fish in group *g* at survey *j* within site *i*, *μ_ijg_* is the expected count, *ϕ_g_* is the dispersion parameter, and *μ_ig_* is the random site effect. Models excluding site-level effects or including zero-inflation performed substantially worse, and estimated zero-inflation probabilities were near zero across all responses. The final model was fitted with four chains of 4000 iterations each (1000 warmup), using control parameters adapt_delta = 0.90 and max_treedepth = 20. Default weakly informative priors were applied, as no custom priors were specified. This included default priors on fixed effects, intercepts, and group-level standard deviations, as implemented in brms (Bürkner 2017). Posterior predictive checks were conducted using pp_check() to confirm adequate model fit. All parameters demonstrated convergence (^R= 1.00), with high effective sample sizes, supporting confidence in the posterior estimates.

Inferences were based on posterior distributions of fixed effects. Predicted mean abundances and proportional group compositions were visualised using faceted bar plots derived from population-level predictions via conditional_effects(). Differences across reef classifications were calculated by combining intercepts with classification-level effects on the log scale, following:

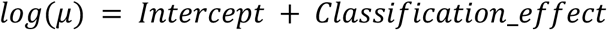

Posterior distributions of fixed effects were extracted using spread_draws() to estimate log-scale differences in functional group abundance across reef classifications. Differences were visualised using stat_halfeye(), which displays the full posterior distribution with overlaid medians and 95% credible intervals.

#### 2.3.2. Zone-based analysis

A separate multivariate model was fitted to examine differences in functional group abundance across geographic zones. This analysis was conducted to assess whether spatial context, specifically, proximity to shore versus offshore exposure, also structured fish assemblages, independent of reef type. The response structure and model family remained identical to the classification-based model, with the predictor variable replaced by zone (nearshore, pelagic, or shipwreck):

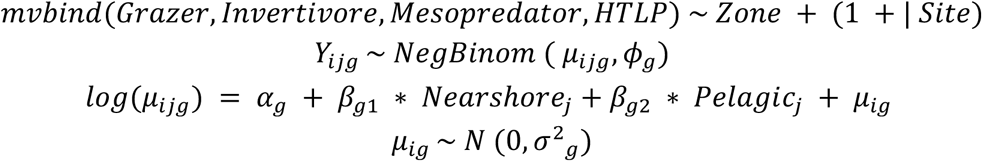

Where *Y_ijg_* represents the count of fish in group *g* at survey *j* within site *i*, *μ_ijg_* is the expected count, *ϕ_g_* is the dispersion parameter, and *μ_ig_* is the random site effect. Shipwrecks were retained as the reference level for interpretability. Model fitting and diagnostics followed the same procedures, including the use of a negative binomial likelihood, site-level random intercepts, and evaluation via LOO-CV. Model diagnostics indicated good convergence and posterior sampling behaviour. LOO-CV showed minimal difference in expected predictive performance compared to the classification model (Δelpd = –0.1), indicating that reef type likely explains functional group composition slightly more effectively than spatial zone. Predicted mean abundances and differences across reef zones were extracted and visualised using the same procedure as the functional group model. Specifically, posterior draws were obtained via spread_draws(), and zone-level effects were visualised using stat_halfeye() with 95% credible intervals, showing log-scale abundance differences relative to shipwrecks.

#### 2.3.3. Species-specific analysis

Species-specific models were fitted to investigate fine-scale differences in abundance across reef classifications. Only species with at least 10 non-zero observations were included to ensure sufficient data for estimation. Survey-level data were aggregated and pivoted to a wide format, and a multivariate negative binomial model was fitted using the brms package, with abundance modelled as a function of reef classification. The response was specified as a multivariate bind of all included species, enabling joint estimation while accounting for overdispersion structured as:

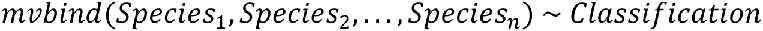

The response for species *s* at survey *j* was modelled as:

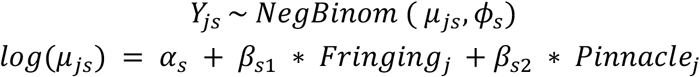

Here, *Y_js_* represents the count of species ss in survey *j*, *μ_js_* is the expected count, *ϕ_s_* is the species-specific dispersion parameter, and shipwrecks were treated as the reference category. Site was not included as a random effect due to limited data and resulting model instability. With 18 taxa across 12 sites, the model became overparameterised and failed to converge reliably. Excluding random intercepts improved stability while retaining the ability to detect habitat-based trends. Model fitting used four chains of 4000 iterations each (1000 warmup) with adapt_delta = 0.90 and max_treedepth = 20. Species-level predictions were generated using conditional_effects() with population-level estimates. Differences in predicted abundance across reef types were visualised using faceted bar plots with 95% credible intervals.

## 3. RESULTS

A total of 350 fish surveys were conducted across 14 unique reef sites, comprising 5 fringing reefs, 6 pelagic pinnacles, and 3 shipwrecks. Repeated surveys were conducted at each site, with 150 surveys on fringing reefs, 130 on pinnacles, and 70 on shipwrecks. Each survey covered an estimated area of ∼1,200 m², yielding a cumulative surveyed area of ∼42 hectares. Across all sites, 162,894 individual fish were recorded. Mean (±SD) fish density was highest on pinnacles (668 ± 928 fish per 1,200 m²), followed by shipwrecks (348 ± 396) and fringing reefs (345 ± 220), indicating elevated average fish abundance at pinnacles and, to a lesser extent, shipwrecks relative to fringing reefs.

### 3.1. Functional groups

Predicted fish abundance varied across reef types and functional groups (Figure 2, Table 1). Grazers and mesopredators were the most abundant groups across all classifications, followed by invertivores and higher trophic level predators (HTLPs). Posterior predictions indicated that grazer abundance was highest at fringing reefs (median: 127.9 fish per 1,200 m²; 95% Credible Interval: [91, 181]), slightly lower at pinnacles (105.9; [78, 145]), and substantially lower at shipwrecks (59.7; [38, 94]). Invertivores showed comparable predicted abundance at pinnacles (56.0; [34, 94]) and fringing reefs (55.9; [32, 97]), with lower estimates at shipwrecks (33.0; [16, 66]). Mesopredator abundance was highest at pinnacles (130.5; [56, 302]), with lower values at fringing reefs (68.7; [26, 172]) and shipwrecks (43.5; [13, 140]). HTLPs had the lowest predicted abundance at fringing reefs (11.0; [5, 23]), with similar values at pinnacles (26.4; [13, 51]) and shipwrecks (25.6; [10, 68]).

**Figure 2.**
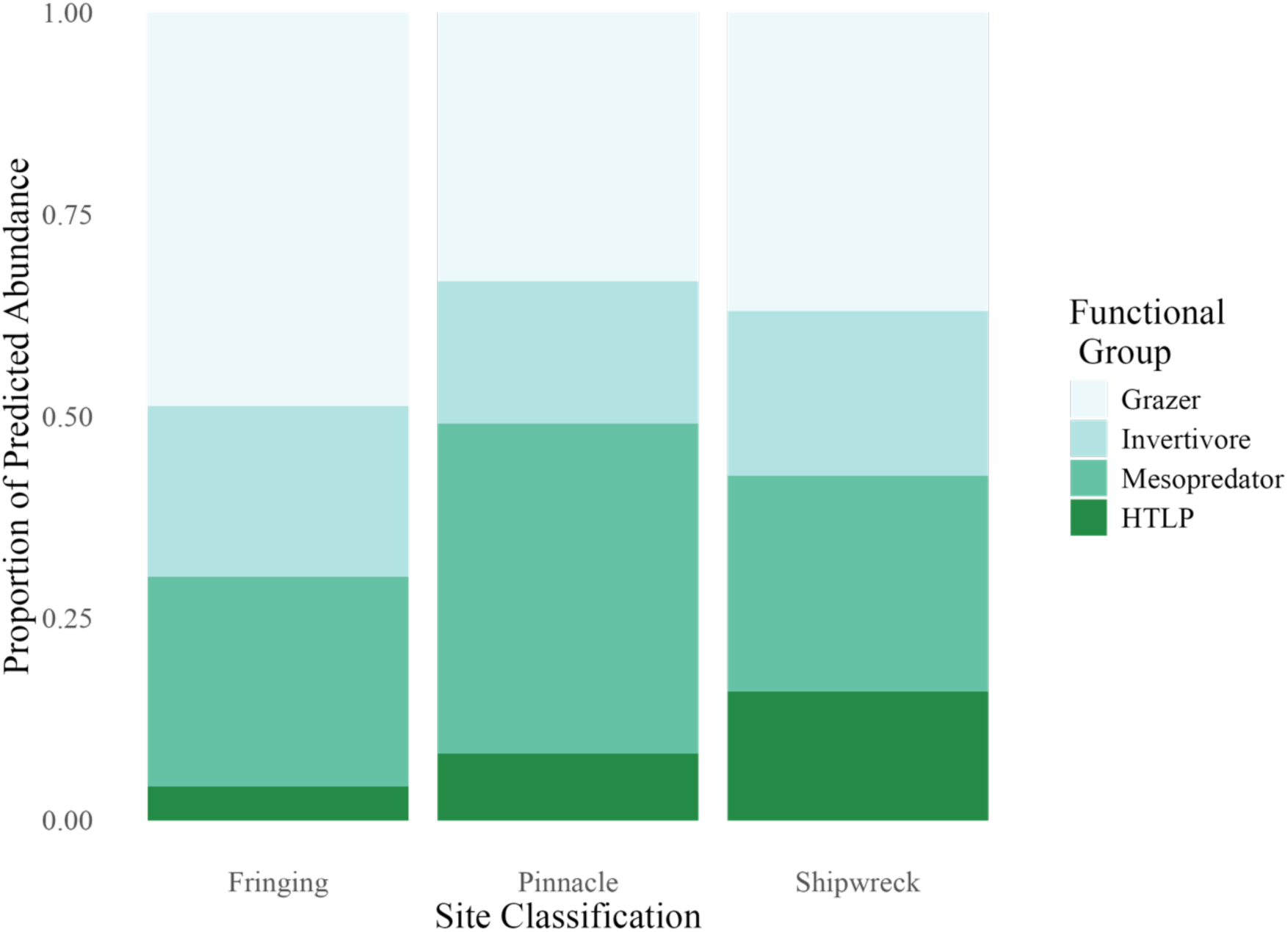
Predicted mean abundances of four functional fish groups—grazers, invertivores, mesopredators, and higher trophic level predators (HTLPs)—across three reef classifications: fringing reefs, pelagic pinnacles, and shipwrecks. Each panel displays the posterior mean abundance for one functional group based on multivariate negative binomial models.

**Table 1.** Predicted mean abundances (with 95% credible intervals) of four functional fish groups across three reef classifications: shipwreck, fringing reef, and pelagic pinnacle. Estimates are derived from a multivariate negative binomial model with reef classification as a fixed effect and site as a random intercept. Values reflect the expected total count per survey for each functional group.

| Functional Group | Shipwreck | Fringing | Pinnacle |
| --- | --- | --- | --- |
| **Grazer** | 59.7 [38.1–94.1] | 127.9 [90.6–180.8] | 105.9 [77.6–145.1] |
| **Invertivore** | 33.0 [15.9–65.8] | 55.9 [31.6–96.7] | 56.0 [33.9–93.6] |
| **Mesopredator** | 43.5 [13.2–140.1] | 68.7 [26.2–171.9] | 130.5 [56.6–301.7] |
| **HTLP** | 25.6 [10.2–68.0] | 11.0 [5.1–23.3] | 26.4 [13.4–51.1] |

Posterior distributions indicated differences in functional group abundance across reef types, with varying levels of statistical support (Figure 3, Table 2). Grazers were more abundant at fringing reefs than shipwrecks, with a median effect of 0.76 (95% CrI: [0.20, 1.34]) and a 99% posterior probability that abundance was higher at fringing reefs. Pinnacles also supported higher grazer abundance (0.57; [0.02, 1.13]), with 98% posterior support for a positive effect. Invertivores followed a similar pattern: abundance was higher at fringing reefs (0.52; [–0.36, 1.42]) and pinnacles (0.52; [–0.34, 1.42]) compared to shipwrecks, with posterior probabilities of 89% and 90%, respectively. Mesopredators were more abundant at pinnacles than shipwrecks (1.10; [–0.35, 2.58]), with 94% support for a positive effect, and may have also been more abundant at fringing reefs (0.45; [–1.07, 1.99]), though posterior support was weaker (74%). For higher trophic level predators (HTLPs), abundance was lower at fringing reefs compared to shipwrecks (–0.86; [–2.10, 0.40]), with a 92% probability of a negative effect. Pinnacles also supported higher HTLP abundance than fringing reefs (0.88; [–0.14, 1.86]), with 96% support. In contrast, the difference between pinnacles and shipwrecks was negligible (0.02; [–1.17, 1.16]), with posterior support evenly split (52%).

**Figure 3.**
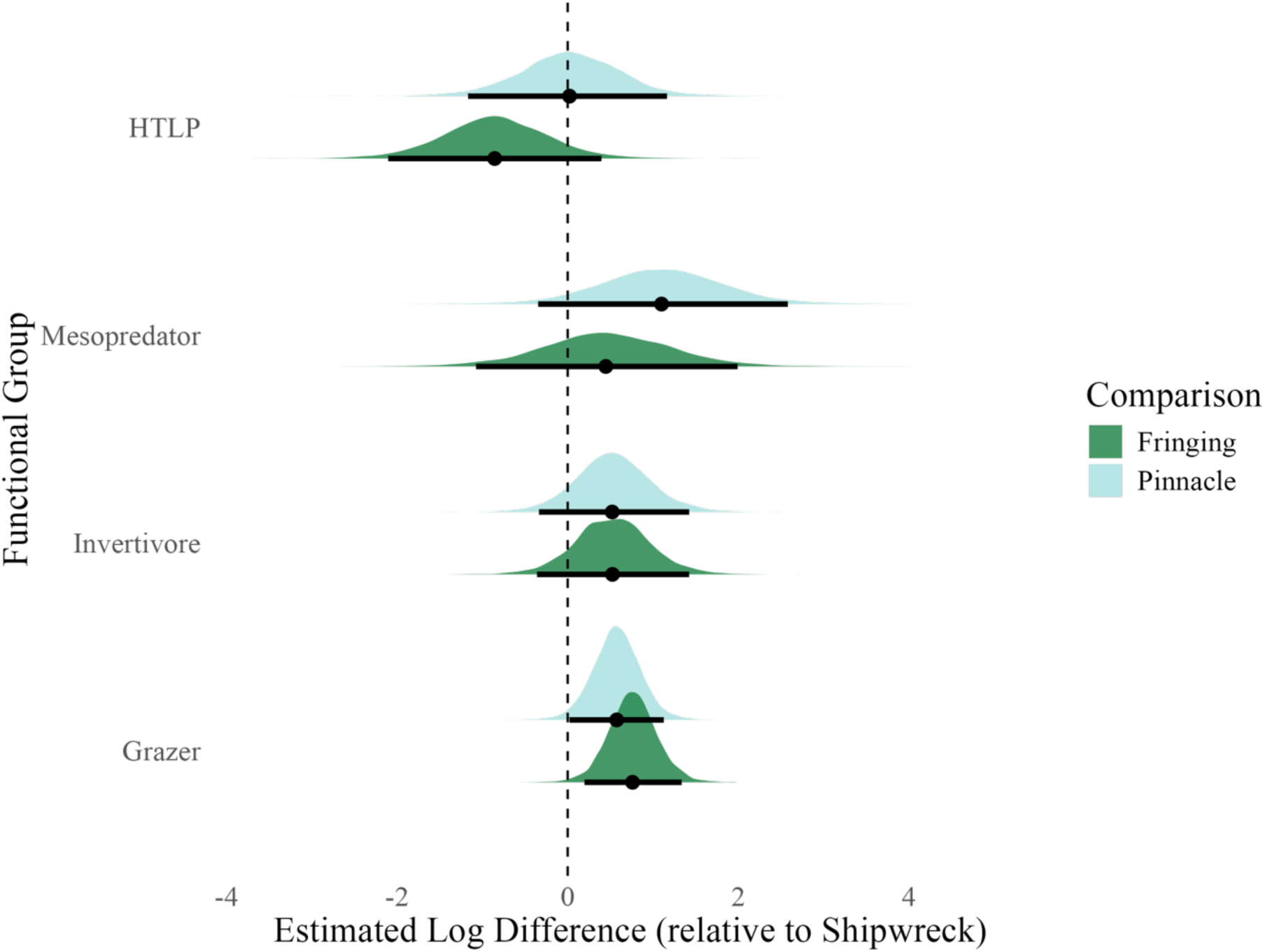
Posterior distributions show estimated log-scale differences in abundance for each functional group on fringing reefs (dark green) and pinnacles (light blue), relative to shipwrecks. Densities represent posterior uncertainty, with medians (dots) and 95% credible intervals (thick bars) overlaid. Zero indicates no difference from shipwrecks; positive values denote higher predicted abundance.

**Table 2.** Posterior summaries of functional group abundance contrasts across zones from the across reef classifications (shipwreck, fringing reef, pinnacle). Values are derived from a Bayesian multivariate negative binomial model. The table reports the posterior median, 95% credible interval (CI), and the proportion of posterior samples supporting positive or negative differences. Estimates are derived from posterior draws of the reef-classification model, with shipwrecks set as the reference level.

| Functional Group | Comparison | Median | 95% CI | P($\beta>0$) | P($\beta<0$) |
| --- | --- | --- | --- | --- | --- |
| **Grazer** | Fringing – Shipwreck | 0.760 | [0.197, 1.340] | 0.994 | 0.006 |
|  | Pinnacle – Shipwreck | 0.574 | [0.022, 1.130] | 0.979 | 0.021 |
|  | Pinnacle – Fringing | –0.188 | [–0.657, 0.278] | 0.203 | 0.797 |
| **Invertivore** | Fringing – Shipwreck | 0.525 | [–0.359, 1.420] | 0.891 | 0.109 |
|  | Pinnacle – Shipwreck | 0.522 | [–0.337, 1.420] | 0.897 | 0.103 |
|  | Pinnacle – Fringing | 0.006 | [–0.733, 0.738] | 0.507 | 0.493 |
| **Mesopredator** | Fringing – Shipwreck | 0.447 | [–1.070, 1.990] | 0.738 | 0.262 |
|  | Pinnacle – Shipwreck | 1.100 | [–0.346, 2.580] | 0.939 | 0.061 |
|  | Pinnacle – Fringing | 0.642 | [–0.616, 1.980] | 0.854 | 0.146 |
| **HTLP** | Fringing – Shipwreck | –0.855 | [–2.100, 0.396] | 0.079 | 0.921 |
|  | Pinnacle – Shipwreck | 0.022 | [–1.170, 1.160] | 0.518 | 0.482 |
|  | Pinnacle – Fringing | 0.880 | [–0.141, 1.860] | 0.960 | 0.041 |

### 3.2. Geographic zones

While reef classification explained most of the variation in functional group composition, a secondary analysis evaluated whether geographic zone, defined by proximity to shore, also structured fish assemblages (Table 3, Figure 4). Grazers were more abundant in both nearshore (median effect: 0.67; 95% CrI: [0.12, 1.28]) and pelagic zones (0.62; [–0.09, 1.32]) compared to shipwrecks, with a 99% and 96% posterior probability, respectively, that abundance was higher in natural zones. Invertivores were also more abundant nearshore (0.67; [–0.08, 1.44]), with a 96% probability that abundance exceeded that on shipwrecks, while the contrast between pelagic sites and shipwrecks was less clear (0.12; [–0.77, 1.05]), supported by a posterior probability of 61%. Mesopredators were somewhat more abundant nearshore (0.84; [–0.68, 2.29]) and in pelagic zones (0.68; [–1.17, 2.49]), with 84% and 78% posterior support, respectively, that abundance was higher than on shipwrecks. For HTLPs, pelagic zones were estimated to support slightly higher abundance than both nearshore reefs (0.57; [–0.68, 1.84]) and shipwrecks (0.07; [–1.45, 1.60]), though posterior support was weak in both cases (57% and 54%, respectively), suggesting no consistent difference in HTLP abundance across zones.

**Figure 4.**
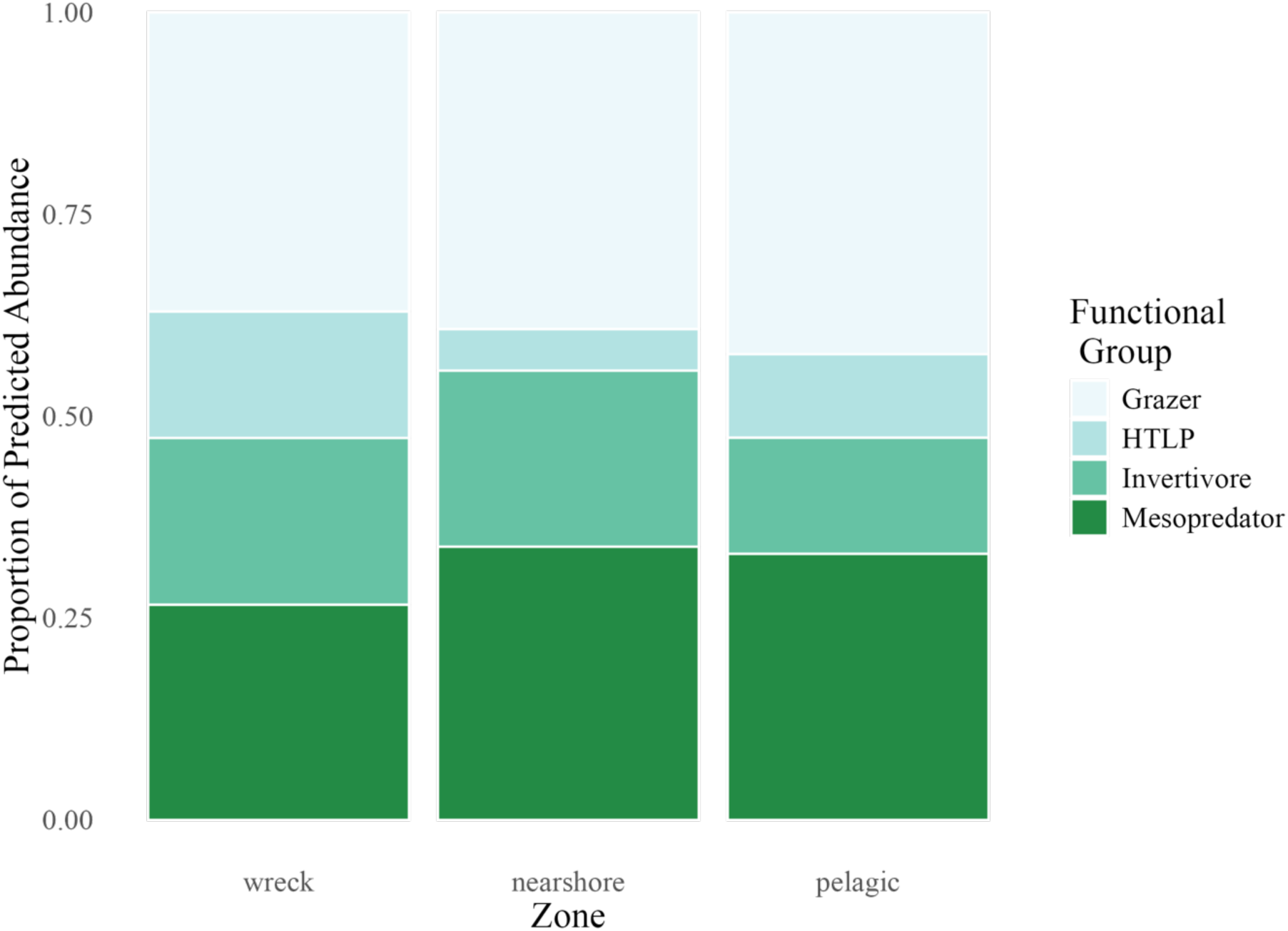
Predicted proportional composition of four functional fish groups across three reef zones: wreck, nearshore, and pelagic. Each bar represents the relative contribution of grazers, invertivores, mesopredators, and higher trophic level predators (HTLPs) to the total fish count within each zone. Proportions are normalised to 1.0 for comparison across zones.

**Figure 5.**
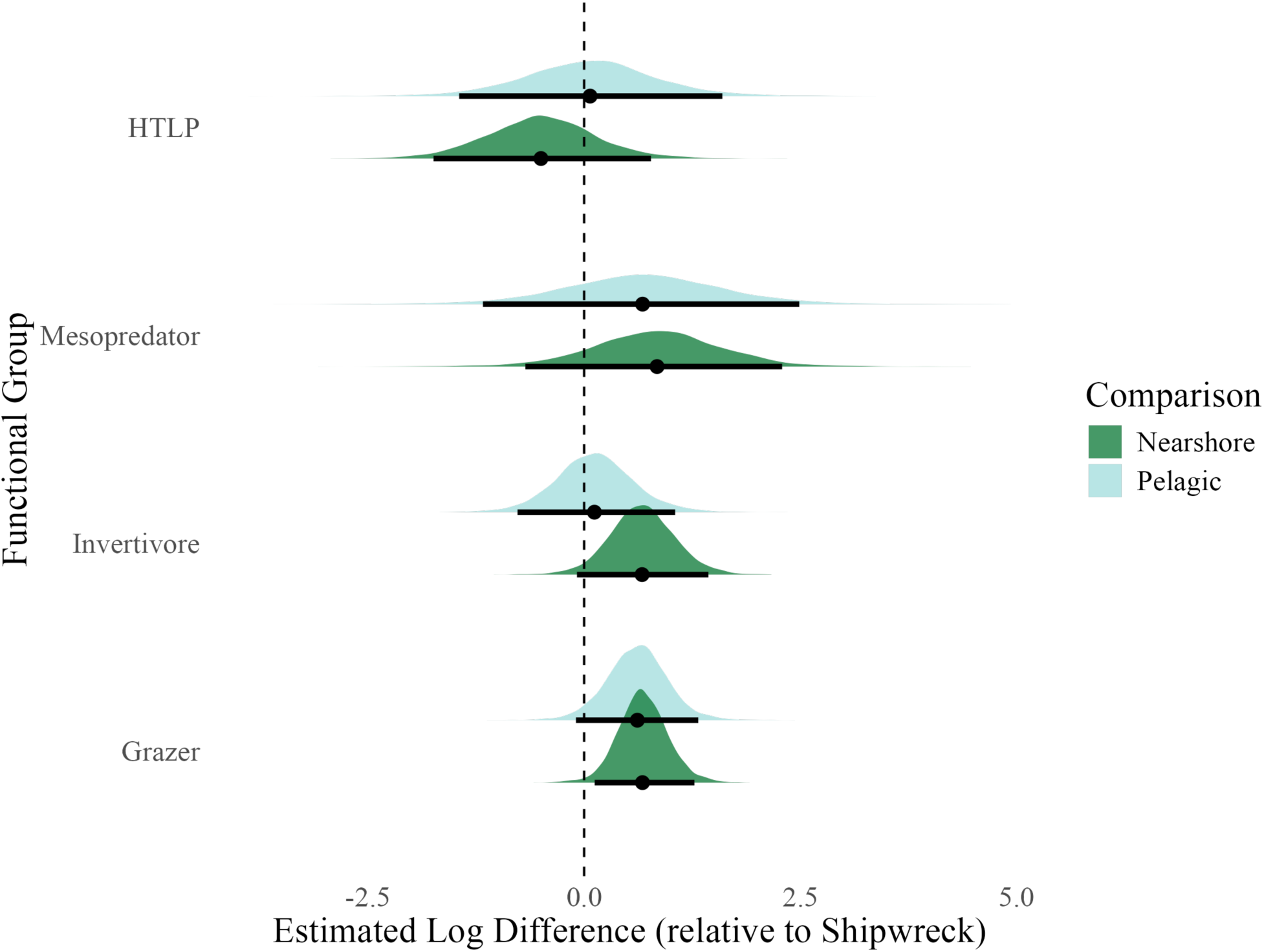
Posterior distributions show estimated log-scale differences in abundance for each functional group in nearshore (dark green) and pelagic (light blue) zones, relative to shipwrecks.Densities represent posterior uncertainty, with medians (dots) and 95% credible intervals (thick bars) overlaid. Zero indicates no difference from shipwrecks; positive values denote higher predicted abundance.

**Table 3.** Posterior summaries of functional group abundance contrasts across zones from the Bayesian multivariate model. The table reports the posterior median, 95% credible interval (CI), and the proportion of posterior samples supporting positive or negative differences. Estimates are derived from posterior draws of the zone-level model, with shipwrecks set as the reference level.

| Functional Group | Comparison | Median | 95% CI | P($\beta > 0$) | P($\beta < 0$) |
| --- | --- | --- | --- | --- | --- |
| **Grazer** | Nearshore – Wreck | 0.674 | [0.122, 1.280] | 0.988 | 0.012 |
|  | Pelagic – Wreck | 0.615 | [−0.095, 1.320] | 0.958 | 0.042 |
|  | Pelagic – Nearshore | −0.066 | [−0.662, 0.505] | 0.404 | 0.596 |
| **Invertivore** | Nearshore – Wreck | 0.670 | [−0.083, 1.440] | 0.964 | 0.036 |
|  | Pelagic – Wreck | 0.119 | [−0.770, 1.050] | 0.606 | 0.394 |
|  | Pelagic – Nearshore | −0.553 | [−1.290, 0.190] | 0.063 | 0.937 |
| **Mesopredator** | Nearshore – Wreck | 0.843 | [−0.679, 2.290] | 0.878 | 0.122 |
|  | Pelagic – Wreck | 0.675 | [−1.170, 2.490] | 0.782 | 0.218 |
|  | Pelagic – Nearshore | −0.163 | [−1.650, 1.340] | 0.408 | 0.592 |
| **HTLP** | Nearshore – Wreck | −0.499 | [−1.740, 0.773] | 0.200 | 0.800 |
|  | Pelagic – Wreck | 0.069 | [−1.450, 1.600] | 0.539 | 0.461 |
|  | Pelagic – Nearshore | 0.569 | [−0.681, 1.840] | 0.819 | 0.181 |

### 3.3. Species-level contrasts

Posterior contrasts for individual species revealed consistent patterns in the direction and strength of habitat associations, particularly distinguishing shipwrecks from natural reef types. Species with the most pronounced or ecologically relevant differences among habitats are highlighted below; full posterior estimates and contrast summaries for all modelled taxa are provided in Tables S2, and S3, and Figures 6–7.

**Figure 6.**
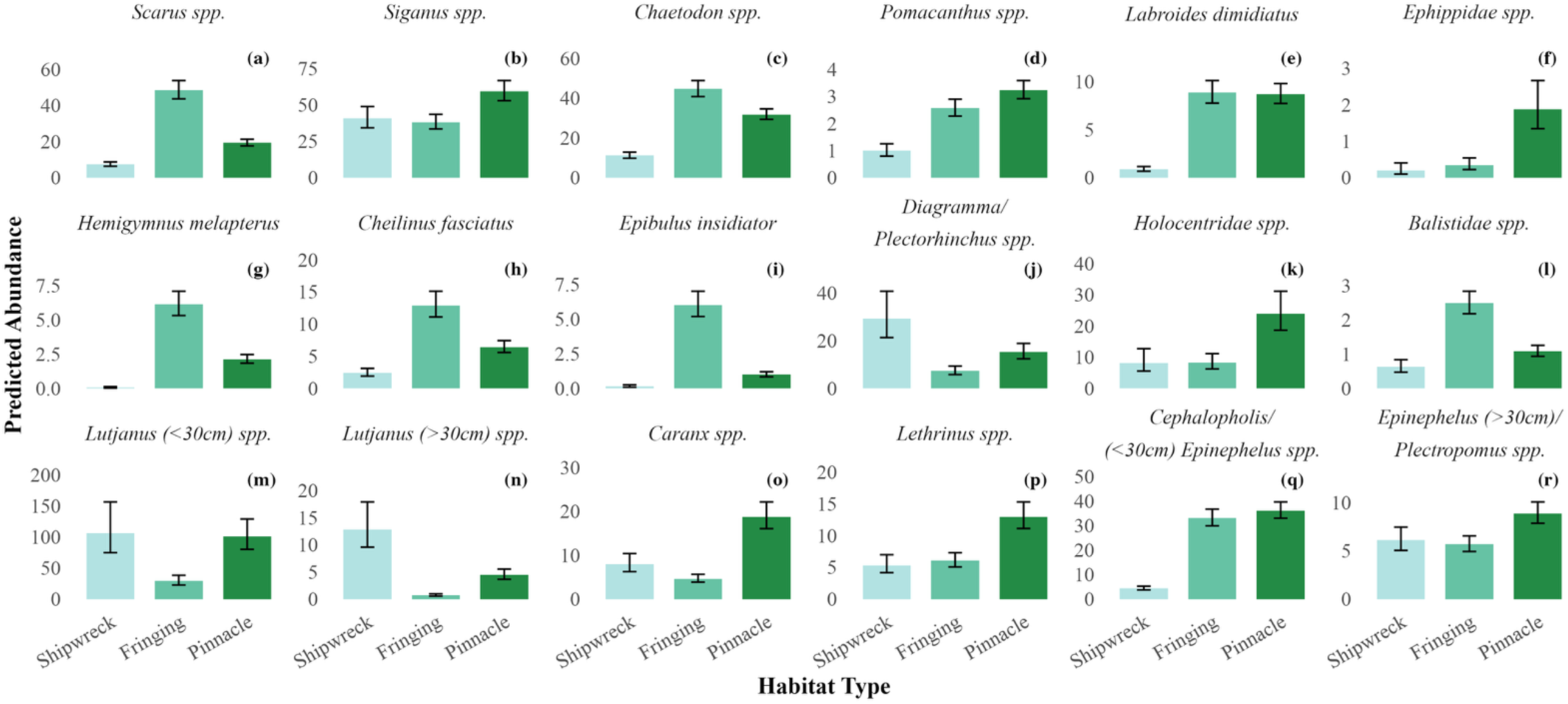
Predicted mean abundances of 18 reef fish taxa (a through r) across three reef classifications: fringing reef, pelagic pinnacle, and shipwreck. Each panel represents one species, with bars showing the posterior mean abundance from multivariate negative binomial models. Error bars represent Bayesian 95% credible intervals. Bars are coloured by reef classification to facilitate comparison of relative abundance patterns across reef types.

**Figure 7.**
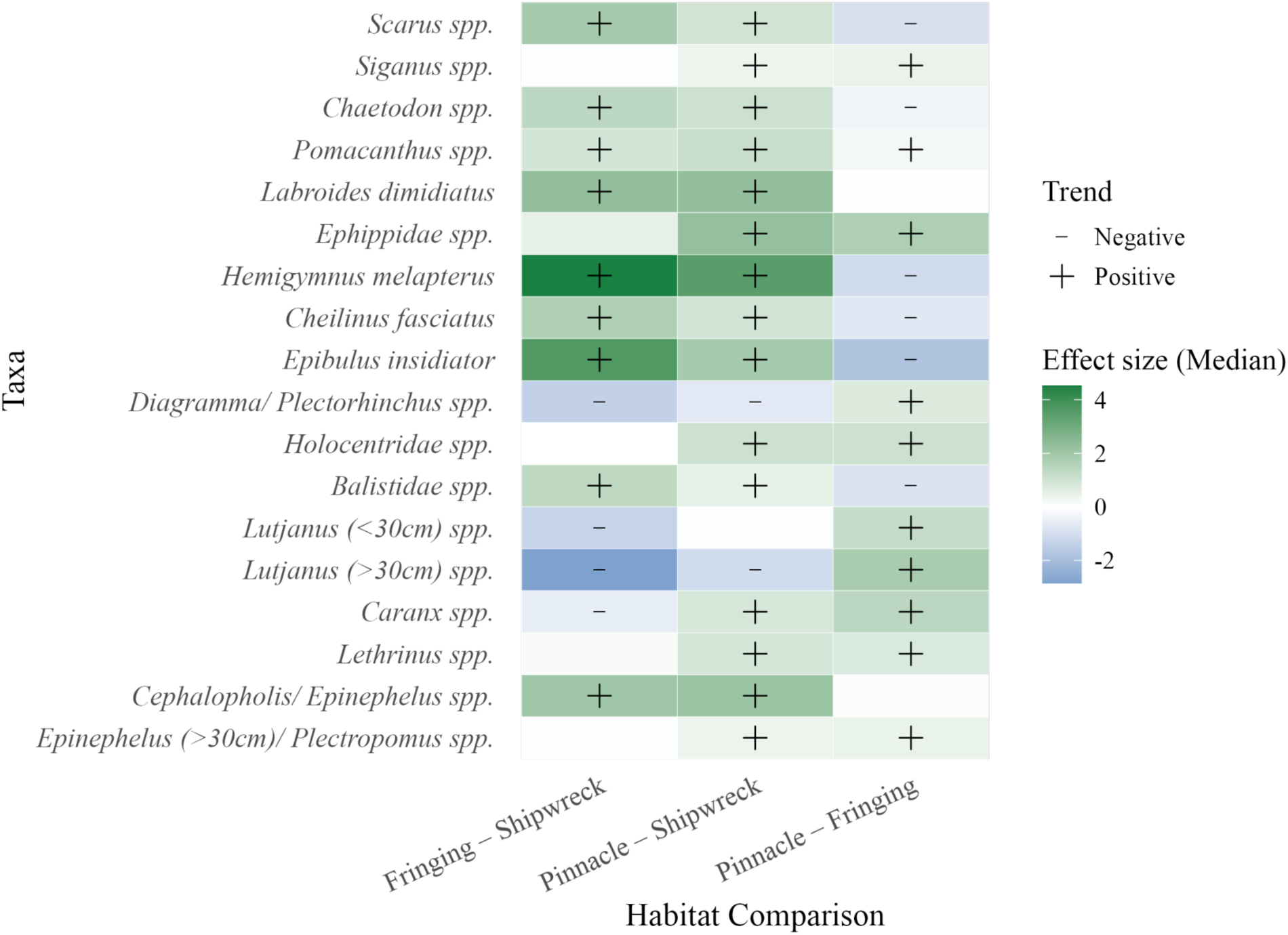
Species-level estimates of fish abundance differences across habitat types, relative to shipwrecks, from a Bayesian multivariate model. Each tile represents the median posterior effect size for a given species–habitat comparison (Fringing or Pinnacle reef vs. Shipwreck baseline). Positive values indicate higher predicted abundance on natural reefs relative to shipwrecks, and negative values indicate the opposite. Black symbols denote species with strong support for directional effects (posterior probability > 0.95 or < 0.05); plus signs indicate positive effects, and minus signs indicate negative effects.

Among mesopredators and high trophic level predators, several species displayed distinct habitat associations. *Lutjanus* spp. <30 cm (small snappers; mesopredators) showed similar predicted abundance at pinnacles and shipwrecks, with a negligible median difference and low posterior support for any effect (–0.05; 95% CrI: [–0.51, 0.38]; 59%), but were substantially less abundant at fringing reefs compared to shipwrecks (–1.29; [–1.75, –0.84]; 100%). *Lutjanus* spp. >30 cm (large snappers; mesopredators) were more abundant at shipwrecks than pinnacles (–1.06; 95% CrI: [–1.44, –0.69]; 100%) and fringing reefs (–2.85; [–3.28, –2.44]; 100%). *Caranx* spp. (trevallies; high trophic level predators) were more abundant at pinnacles compared to shipwrecks (0.85; 95% CrI: [0.55, 1.15]; 100%) and more abundant at shipwrecks relative to fringing reefs (–0.54; [–0.86, –0.23]; 100%). *Diagramma/ Plectorhinchus* spp. (sweetlips; invertivores) were less abundant at both fringing reefs (–1.38; 95% CrI: [–1.79, –0.98]; 100%) and pinnacles (–0.65; [–1.05, –0.27]; 100%) compared to shipwrecks. *Lethrinus* spp. (emperors; mesopredators) showed substantially higher predicted abundance at pinnacles relative to shipwrecks (0.89; 95% CrI: [0.57, 1.19]; 100%), while differences between fringing reefs and shipwrecks were minor and weakly supported (0.13; [–0.20, 0.44]; 78%). *Epinephelus/ Plectropomus* spp. >30cm (large groupers; high trophic level predators) were more abundant at pinnacles relative to shipwrecks (0.37; 95% CrI: [0.14, 0.60]; 100%), with slightly higher but weakly supported differences at fringing reefs compared to shipwrecks (–0.07; [–0.32, 0.16]; 72%).

In contrast, several grazer and invertivore taxa were consistently lower on shipwrecks relative to natural reefs. *Pomacanthus spp.* (angelfish; invertivores) showed higher predicted abundance at both fringing reefs (0.94; 95% CrI: [0.69, 1.20]; 100%) and pinnacles (1.17; [0.92, 1.41]; 100%) compared to shipwrecks, with minimally elevated abundance at pinnacles relative to fringing reefs (0.23; [0.07, 0.39]; 99.8%). *Labroides dimidiatus* (cleaner wrasse; invertivores) also had higher predicted abundance at fringing reefs (2.29; 95% CrI: [1.99, 2.59]; 100%) and pinnacles (2.27; [1.98, 2.57]; 100%) compared to shipwrecks. *Epibulus insidiator* (slingjaw wrasse; invertivores) showed a similar pattern, with higher abundance at fringing reefs (3.64; [3.12, 4.24]; 100%) and pinnacles (1.86; [1.33, 2.46]; 100%) relative to shipwrecks. *Scarus* spp. (parrotfish; grazers) had higher predicted abundance at fringing reefs (1.87; [1.68, 2.06]; 100%) and pinnacles (0.95; [0.77, 1.14]; 100%) compared to shipwrecks. *Chaetodon* spp. (butterflyfish; grazers) followed a similar pattern, with elevated abundance at fringing reefs (1.35; [1.06, 1.64]; 100%) and pinnacles (0.65; [0.36, 0.94]; 100%) relative to shipwrecks. Unlike other grazer taxa, *Siganus* spp. (rabbitfish; grazers) exhibited minimal variation, with slightly higher abundance on pinnacles (0.37; [0.15, 0.58]; 100%) and no strong difference between fringing reefs and shipwrecks (–0.07; [–0.29, 0.15]; 72% probability of a negative effect).

Across species, predatory taxa often reached comparable or higher abundance on shipwrecks compared with pinnacles, whereas reef-specialist grazers and invertivores were consistently lower on artificial structures (Figure 7). The consistently lower abundance of key grazers and invertivores indicates that meaningful compositional differences persist between artificial and natural reef assemblages.

## 4. DISCUSSION

This study evaluated fish assemblage structure across shipwrecks, pinnacles, and fringing reef environments to determine whether artificial reefs replicate natural community patterns. Based on their vertical relief and relative isolation, shipwrecks were expected to support assemblages more similar to pinnacles than found on fringing reefs. By modelling both functional group and species-level patterns, results suggest that shipwreck assemblages closely resembled natural reef pinnacles in their support of mesopredators and higher trophic level predators, but diverged markedly in their low representation of grazers and invertivores. While several predatory and mobile species were abundant on both pinnacles and shipwrecks, many wrasses, parrotfishes, and small groupers were consistently underrepresented on artificial structures. These patterns highlight the functional selectivity of artificial reefs and their potential to replicate certain ecological roles while failing to support others, reinforcing the need for functionally driven artificial reef designs and deployment strategies, and for natural reef conservation to remain a priority in Thailand (Baillie et al. 2024; Paxton et al. 2020a). Key outcomes are examined in terms of predator convergence, divergence in lower trophic guilds, and implications for artificial reef design and ecological function.

### 4.1. Shipwrecks resemble pinnacles in meso- and higher trophic level predators

Predator-dominated assemblages on shipwrecks closely mirrored those observed on pelagic pinnacles, aligning with the initial prediction. High trophic level predator (HTLP) abundance was particularly elevated on shipwrecks, and larger predators were consistently more abundant at both shipwreck and pinnacle sites than on fringing reefs. This convergence is likely driven by shared physical features: vertical relief, isolation, and strong hydrodynamic flow are known to promote predator aggregation on both pinnacles and artificial reefs (Rilov & Benayahu 2000, Galbraith 2021, Galbraith et al. 2023, Cresswell et al. 2023). Vertical extent has also been identified as a key predictor of predator density on artificial structures, including shipwrecks and oil platforms (Paxton et al. 2020b, Harvey et al. 2021, Sibley et al. 2023, Schram et al. 2024). These features may enhance foraging efficiency and attract mobile predators from surrounding areas, potentially having implications for fish diversity at adjacent reef sites.

These findings are consistent with broader patterns observed across artificial reef systems, which frequently support dense assemblages of schooling pelagic and mobile mid-to-high trophic level predators (Streich et al. 2017, Schulze et al. 2020, Sibley et al. 2023). In this study, predators such as *Lutjanus, Epinephelus / Plectropomus, Caranx,* and *Diagramma / Plectorhinchus* spp. were particularly abundant on shipwrecks, echoing patterns reported in the Gulf of Thailand (Harvey et al. 2021, Baillie et al. 2024). In contrast, Paxton et al. (2020b) found that artificial reefs in the USA supported higher densities of transient species such as *Caranx*, but not resident predators like *Lutjanus* or *Epinephelus*, highlighting potential regional differences in species responses to reef structure. The strong resemblance between shipwreck and pinnacle predator assemblages observed in the current study suggests that ship-based artificial reefs may function more like pinnacle reef systems than fringing reefs in terms of predatory fish structure, potentially providing vital insights into management decision making for artificial reef deployments in the region.

### 4.2. Divergence in lower trophic guilds

Despite strong similarities in predator assemblages, shipwrecks diverged markedly from both natural reef types in their support of lower trophic guilds. Grazers and invertivores were consistently underrepresented, with higher densities of *Scarus*, *Chaetodon*, *Pomacanthus*, and various labrids (wrasses) observed on fringing and pinnacle reefs. These patterns appear consistent with broader trends across tropical artificial reef systems, where small-bodied benthic feeders and cryptic invertivores are often scarce (Mattos & Yeemin 2020, Koeck et al. 2014, Hickman et al. 2024, Schram et al. 2024). Notably, *Labroides* and *Hemigymnus* spp. have also been reported as underrepresented on shipwrecks relative to natural pinnacles within the same region (Baillie et al. 2024), reinforcing the likelihood that artificial reefs tend to support distinct assemblages lacking key functional groups associated with benthic foraging. These patterns suggest a degree of functional divergence from natural reef communities, driven by limitations in habitat complexity and resource availability on artificial structures (Fowler & Booth 2012, Simon et al. 2013, Schram et al. 2024).

Several ecological and physical constraints likely contribute to this divergence. Structural complexity is likely insufficient to establish shipwrecks as effective ecological analogues of natural reefs; shipwrecks often lack fine-scale refugia, cryptic microhabitats, and turf algal cover needed by benthic feeders (Simon et al. 2013, Koeck et al. 2014, Mattos & Yeemin 2020, Monchanin et al. 2021, Monchanin et al. 2025), Differences in depth, location, and reef structure may also play a role. However, the nearby pinnacles surveyed at similar depths supported higher densities of grazers and invertivores. Zone-based analyses confirmed that trophic divergence persisted, suggesting that physical setting alone cannot fully explain the consistent underrepresentation of lower trophic groups on shipwrecks. Strong currents around wrecks may further limit benthic foraging and favour species that feed higher in the water column (Galbraith 2021).

*Siganus* spp. were notable for their consistent abundance across reef types, diverging from the patterns observed for other grazer taxa. This may reflect species-level variation in ecological traits. Some of the *Siganus* species recorded in this study primarily feed on turf algae, but others surveyed, such as *Siganus javus*, are known to exhibit more flexible or planktivorous feeding strategies (Streit et al. 2015), potentially allowing persistence on shipwrecks where benthic forage is limited. Their presence on structurally complex artificial reefs has also been reported in other parts of Thailand (Mattos & Yeemin 2020), suggesting a degree of dietary or habitat plasticity not shared by other herbivorous taxa. Additionally, a dominance of planktivores has been observed on shipwrecks in other regions, including the eastern Mediterranean (Sensurat-Genc et al. 2022), suggesting that the structural features of wrecks may consistently favour planktivorous species. These patterns highlight that most benthic-feeding species remain functionally excluded from shipwreck communities unless habitat and resource limitations are addressed explicitly in artificial reef design.

### 4.3. Functional limitations of ARs as ecosystem surrogates

While overall fish density on shipwrecks was high, comparable to fringing reefs and second only to pinnacles, it was driven primarily by mesopredators and higher trophic level predators, with grazers and invertivores consistently underrepresented. Similar patterns have been documented elsewhere: artificial reefs often match or exceed natural reefs in total fish abundance but differ markedly in functional composition, with consistently reduced species diversity/ richness (Fowler & Booth, 2012, Paxton et al. 2020). Assemblages dominated by predators and lacking lower trophic groups may reflect a combination of habitat features, species-specific responses, and top-down effects, where predator presence suppresses prey guilds (Boaden and Kingsford, 2015). Artificial reef fish communities are thus compositionally novel and shaped by deployment context, structural design, and regional species pools (Folpp et al. 2013, Schulze et al. 2020). Coral surveys at the same reef sites included in this study found that communities on artificial structures remain distinct from those on natural reefs, showing partial convergence with pinnacle assemblages over time (Monchanin et al. 2021, Monchanin et al. 2025). While shipwrecks may offer fisheries value and act as de facto refuges under high exploitation pressure as they serve as physical barriers to bottom-trawling practices (Hickman et al. 2024), they appear to fall short of replicating the full ecological function of natural reefs. These findings underscore the need for functionally driven reef design and context-aware management goals when deploying artificial structures.

### 4.4. Design and management implications

Recognising these structural and ecological mismatches is essential for guiding the design of artificial reefs that support a broader range of functional groups. Artificial reefs can play a valuable role in marine conservation and fisheries management (Fowler & Booth 2012, Paxton et al. 2020a, Sibley et al. 2023), but their ecological function depends heavily on how they are designed and where they are deployed (Streich et al. 2017, Paxton et al. 2020a, Baillie et al. 2024). Structures that offer vertical relief are highly effective at attracting predators, particularly mobile and pelagic species (Simon et al. 2013, Paxton et al. 2020b). However, replicating the full ecological breadth of natural reefs requires more than height alone (Monchanin et al. 2021, Schram et al. 2024, Monchanin et al. 2025). Small grazers and invertivores depend on benthic complexity: fine-scale refugia, branching substrates, crevices, and textured surfaces that support turf algae and invertebrate prey; features that are often lacking from artificial reef structures (Koeck et al. 2014; Mattos & Yeemin 2020). Studies show that modular units and internally complex structures can promote the colonisation of benthic specialists more effectively than uniform concrete blocks or ship hulls (Monchanin et al. 2021, Monchanin et al. 2025, Jaxion-Harm et al. 2018, Baillie et al. 2024). Internal cavities and varied surface materials that shipwrecks may lack, can help to improve habitat heterogeneity and support functional diversity in fish assemblages.

The surrounding environment may also strongly shape reef performance. Hydrodynamic conditions, such as current strength and direction, influence the foraging efficiency of pelagic feeders and may enhance the persistence of planktivores and filter feeders (Galbraith et al. 2023, Mattos et al. 2023). Site context matters as much as structure: while deploying artificial reefs near natural reefs may enhance coral recruitment (Monchanin et al. 2021, Monchanin et al. 2025), it may not necessarily ensure ecological similarity or functional equivalence in fish assemblages (Simon et al. 2013, Galbraith, 2021). Emerging evidence suggests that artificial reefs placed near natural reefs may displace fish and fragment populations rather than enhance overall biodiversity (Layman & Allegier 2020; Medeiros et al. 2022, Paxton et al. 2023). This is central to the attractor versus producer hypothesis that is subject to debate in the artificial reef literature (see Paxton et al. 2023 review). Such displacement can also increase vulnerability to exploitation, particularly when artificial habitats draw fish away from protected areas. However, other studies report limited effects of proximity to natural reefs on fish assemblages at shipwrecks (Ross et al. 2016, Sensurat-Genc et al. 2022). Even where displacement occurs, it may not be ecologically detrimental in all contexts. In areas with high fishing pressure, shipwrecks can act as de facto marine refuges, supporting greater fish biomass and more structured communities than surrounding trawled zones (Hickman et al. 2024). These findings highlight the importance of spatial planning that prioritises functional complementarity with natural reefs, while also recognising the potential of ARs to provide refuge and mitigate anthropogenic impacts in degraded seascapes.

Beyond site-level design, broader management strategies in the Gulf of Thailand should ensure that artificial reefs are paired with natural reef conservation to ensure ecosystem function is maintained. While shipwrecks and similar structures can support fisheries management, particularly with larger predators (Harvey et al. 2021), they are not ecological replacements for natural reefs and must be integrated within broader reef management strategies that prioritise ecosystem conservation (Streich et al. 2017, Sibley et al. 2023). Furthermore, given that nearshore shipwrecks have been associated with reduced community heterogeneity and limited ecological benefits (Medeiros et al. 2022), offshore deployments may offer greater functional value. Positioning artificial reefs in offshore settings that mimic the vertical structure of natural pinnacles could enhance support for vulnerable and commercially important species. Achieving trophic balance requires purpose-built artificial reefs designed to support a range of functional groups, not just HTLP species (Paxton et al. 2020a). Structures should be tailored to ecological objectives, incorporating habitat complexity for lower trophic guilds and sited based on hydrodynamic conditions and connectivity. Evaluations must go beyond fish abundance to consider trophic structure, mutualist presence, and long-term community composition. As artificial reefs tend to form novel assemblages (Fowler & Booth 2012, Folpp et al. 2013) functional outcomes, not just biomass metrics, should guide design, deployment, and monitoring (Paxton et al. 2020a, Baillie et al. 2024, Scram et al. 2024). Management frameworks and permitting processes should reflect this by linking artificial reef approval to clearly defined ecological functions and ensuring alignment with regional reef conservation priorities. Ultimately, artificial reefs have the potential to complement conservation goals, but protecting and restoring natural reefs remains essential to sustaining biodiversity and ecosystem resilience in the region.

Where this study focused on variance of fish assemblages across natural and artificial reef sites, future research should extend beyond fish to include benthic and fouling communities, which influence habitat development and substrate use over time (Fowler & Booth 2012). Long-term monitoring of colonisation and successional trajectories as done in Monchanin et al. (2021, 2025) will help capture shifts in community composition and ecosystem function (Folpp et al. 2013). Structural comparisons between modular and monolithic designs, particularly with respect to internal complexity, may reveal strategies for enhancing habitat suitability for lower trophic guilds (Mattos & Yeemin 2020). Incorporating hydrodynamic modelling could improve understanding of spatial patterns in fish abundance and diversity (Galbraith et al. 2023), while behavioural tracking tools, such as acoustic telemetry, can clarify patterns of fish movement, site fidelity, and habitat use. Seasonal monitoring and assessments of invasive species and habitat connectivity will also be critical for managing long-term reef health (Jaxion-Harm et al. 2018, Schulze et al. 2020). Overall, this study provides a foundation for evaluating functional performance and outcomes of artificial reefs. Ongoing research should prioritise context-specific design, ecological function, and long-term monitoring to inform adaptive management and reef policy in tropical systems. Together, these efforts will help ensure that artificial reefs contribute meaningfully to ecosystem resilience, biodiversity conservation, and sustainable reef management in the face of accelerating environmental change.

## 5. CONCLUSION

This study is the first to compare shipwrecks, pelagic pinnacles, and fringing reefs in terms of fish assemblage structure. Shipwrecks mirrored pinnacles in supporting mesopredators and higher trophic level predators but consistently lacked grazers and invertivores compared to both natural reef types, resulting in a skewed trophic balance. These patterns likely reflect structural and ecological limitations, though site-level variability and seasonal effects may also play a role. Future research using long-term monitoring, fine-scale habitat surveys, and behavioural data will help clarify the drivers of these differences. Effective deployment depends on more than high relief and isolation. Designs should incorporate fine-scale complexity to support lower trophic guilds, and site selection should consider hydrodynamics, connectivity, and spatial placement relative to natural reefs. In Thailand and other tropical regions, high-relief offshore structures may better meet objectives for vulnerable and commercially important species. Artificial reefs can complement, but not replace, the protection and restoration of natural reefs, and their success depends on clear ecological goals, functional design, and integration within broader conservation strategies.

## 8. CRediT AUTHOR CONTRIBUTIONS

**Scarlett R. Taylor:** Methodology, Investigation, Software, Formal Analysis, Visualisation, Writing - Original draft preparation, Writing - Reviewing and Editing. **Gavin Miller:** Conceptualisation, Methodology, Investigation, Writing - Original draft preparation, Writing - Reviewing and Editing. **Piers Baillie:** Methodology, Investigation, Project Administration, Supervision, Writing - Reviewing and Editing.

## 9. ACKNOWLEDGEMENTS

The authors thank all Global Reef interns and research assistants who contributed to data collection during the study period, and Hydronauts Diving Resort for their excellent logistical support. We also acknowledge the DMCR for their ongoing support of marine research in Thailand.

## 12. ETHICAL APPROVAL

This study did not require formal ethical approval, and data collection followed standard scientific ethical guidelines. All necessary permits for observational research were secured.

## 13. DATA AVAILABILITY

All data and code used in this study are openly available through Zenodo from https://doi.org/10.5281/zenodo.16784464, archived at the time of submission.

## REFERENCES

Baillie, P., Miller, G., and Chaona, G. (2024). Assessing the performance of Artificial Reefs as a fisheries management tool in the Gulf of Thailand. Phuket Marine Biological Center Research Bulletin, 81, 75–90. 10.14456/pmbcrb.2024.4

Bell, J. D., and Galzin, R. (1984). Influence of live coral cover on coral-reef fish communities. Marine Ecology Progress Series, 15, 265–274. https://www.int-res.com/articles/meps/15/m015p265.pdf

Boaden, A. E., and Kingsford, M. J. (2015). Predators drive community structure in coral reef fish assemblages. In Ecosphere (Vol. 6, Issue 4). Ecological Society of America. 10.1890/ES14-00292.1

Bürkner, P.-C. (2017). brms: An R package for Bayesian multilevel models using Stan. Journal of Statistical Software, 80(1), 1–28. 10.18637/jss.v080.i01

Cresswell, B., Galbraith, G., Harrison, H., McCormick, M., and Jones, G. (2023). Coral reef pinnacles act as ecological magnets for the abundance, diversity and biomass of predatory fishes. Marine Ecology Progress Series, 717, 143–156. 10.3354/meps14377

Folpp, H., Lowry, M., Gregson, M., and Suthers, I. M. (2013). Fish Assemblages on Estuarine Artificial Reefs: Natural Rocky-Reef Mimics or Discrete Assemblages? PLoS ONE, 8(6). 10.1371/journal.pone.0063505

Fowler, A. M., and Booth, D. J. (2012). How well do sunken vessels approximate fish assemblages on coral reefs? Conservation implications of vessel-reef deployments. Marine Biology, 159(12), 2787–2796. 10.1007/s00227-012-2039-x

Galbraith, G. (2021). The ecology of fishes on submerged pinnacle coral reefs. PhD Thesis, James Cook University. 10.25903/7ek9%2D5m80

Galbraith, G., Mcclure, E. C., and Barnett, A. (2022). Diving into the Deep: The Unique Deep Habitats of the Coral Sea Marine Park. Report prepared for Parks Australia. 10.13140/RG.2.2.10516.68487/1

Galbraith, G. F., Cresswell, B. J., McCormick, M. I., and Jones, G. P. (2023). Strong hydrodynamic drivers of coral reef fish biodiversity on submerged pinnacle coral reefs. Limnology and Oceanography, 68(11), 2415–2430. 10.1002/lno.12431

Harvey, E. S., Watts, S. L., Saunders, B. J., Driessen, D., Fullwood, L. A. F., Bunce, M., Songploy, S., Kettratad, J., Sitaworawet, P., Chaiyakul, S., Elsdon, T. S., and Marnane, M. J. (2021). Fish Assemblages Associated With Oil and Gas Platforms in the Gulf of Thailand. Frontiers in Marine Science, 8, 664014. 10.3389/FMARS.2021.664014/BIBTEX

Hickman, J., Richards, J., Rees, A., and Sheehan, E. v. (2024). Shipwrecks act as de facto Marine Protected Areas in areas of heavy fishing pressure. Marine Ecology, 45(1). 10.1111/maec.12782

Jaxion-Harm, J., Szedlmayer, S. T., and Mudrak, P. A. (2018). A Comparison of Fish Assemblages According to Artificial Reef Attributes and Seasons in the Northern Gulf of Mexico. In American Fisheries Society Symposium (Vol. 86). https://sedarweb.org/documents/sedar-74-rd09-a-comparison-of-fish-assemblages-according-to-artificial-reef-attributes-and-seasons-in-the-northern-gulf-of-mexico/

Koeck, B., Tessier, A., Brind’Amour, A., Pastor, J., Bijaoui, B., Dalias, N., Astruch, P., Saragoni, G., and Lenfant, P. (2014). Functional differences between fish communities on artificial and natural reefs: a case study along the French Catalan coast. Aquatic Biology, 20(3), 219–234. 10.3354/ab00561

Layman, C. A., & Allgeier, J. E. (2020). An ecosystem ecology perspective on artificial reef production | Enhanced Reader. Journal of Applied Ecology, 57, 2139–2148. 10.1111/1365-2664.13748

Letessier, T. B., Mouillot, D., Bouchet, P. J., Vigliola, L., Fernandes, M. C., Thompson, C., Boussarie, G., Turner, J., Juhel, J. B., Maire, E., Julian Caley, M., Koldewey, H. J., Friedlander, A., Sala, E., and Meeuwig, J. J. (2019). Remote reefs and seamounts are the last refuges for marine predators across the Indo-Pacific. PLOS Biology, 17(8), e3000366. 10.1371/JOURNAL.PBIO.3000366

Mattos, F. M. G., and Yeemin, T. (2020). First Insight on the Fish Species Composition at Two Artificial Reef Areas, Narathiwat Province, Thailand. Ramkhamhaeng International Journal of Science and Technology, 3(2), 16–23. https://ph02.tci-thaijo.org/index.php/RIST/article/view/193777

Medeiros, A. P. M., Ferreira, B. P., Betancur-R, R., Cardoso, A. P. L. R., Matos, M. R. S. B. C., and Santos, B. A. (2022). Centenary shipwrecks reveal the limits of artificial habitats in protecting regional reef fish diversity. Journal of Applied Ecology, 59(1), 286–299. 10.1111/1365-2664.14053

Monchanin, C., Desmolles, M., and Mehrotra, R. (2025). Homogenization and distinction of coral recruit communities between natural and artificial substrates at Koh Tao a decade after deployment. Aquatic Ecology, 59(2), 597–608. 10.1007/s10452-025-10182-1

Monchanin, C., Mehrotra, R., Haskin, E., Scott, C. M., Urgell Plaza, P., Allchurch, A., Arnold, S., Magson, K., and Hoeksema, B. W. (2021). Contrasting coral community structures between natural and artificial substrates at Koh Tao, Gulf of Thailand. Marine Environmental Research, 172. 10.1016/j.marenvres.2021.105505

Paxton, A. B., Shertzer, K. W., Bacheler, N. M., Kellison, G. T., Riley, K. L., and Taylor, J. C. (2020a). Meta-analysis reveals artificial reefs can be effective tools for fish community enhancement but are not one-size-fits-all. Frontiers in Marine Science, 7. 10.3389/fmars.2020.00282

Paxton, A. B., Newton, E. A., Adler, A. M., van Hoeck, R. v., Iversen, E. S., Christopher Taylor, J., Peterson, C. H., and Silliman, B. R. (2020b). Artificial habitats host elevated densities of large reef-associated predators. PLoS ONE, *15*(9 September). 10.1371/journal.pone.0237374

Paxton, A. B., McGonigle, C., Damour, M., Holly, G., Caporaso, A., Campbell, P. B., Meyer-Kaiser, K. S., Hamdan, L. J., Mires, C. H., & Taylor, J. C. (2023). Shipwreck ecology: Understanding the function and processes from microbes to megafauna. BioScience, 74(1), 12–24. 10.1093/BIOSCI/BIAD084

Rilov, G., and Benayahu, Y. (2000). Fish assemblage on natural versus vertical artificial reefs: The rehabilitation perspective. Marine Biology, 136(5), 931–942. 10.1007/S002279900250

Ross, S. W., Rhode, M., Viada, S. T., & Mather, R. (2016). Fish species associated with shipwreck and natural hard-bottom habitats from the middle to outer continental shelf of the Middle Atlantic Bight near Norfolk Canyon. Fishery Bulletin, 114(1), 45–57. 10.7755/FB.114.1.4

Schram, M. J., Emory, M. E., Kilborn, J. P., Peake, J. A., Wall, K. R., Williams, I., and Stallings, C. D. (2024). Reef fish assemblages differ both compositionally and functionally on artificial and natural reefs in the eastern Gulf of Mexico. ICES Journal of Marine Science, 81(6), 1150–1163. 10.1093/icesjms/fsae075

Schulze, A., Erdner, D. L., Grimes, C. J., Holstein, D. M., and Miglietta, M. P. (2020). Artificial Reefs in the Northern Gulf of Mexico: Community Ecology Amid the “Ocean Sprawl.” In Frontiers in Marine Science (Vol. 7). Frontiers Media S.A. 10.3389/fmars.2020.00447

Şensurat-Genç, T., Lök, A., Özgül, A., and Oruç, A. Ç. (2022). No effect of nearby natural reef existence on fish assemblages at shipwrecks in the Aegean Sea. Journal of the Marine Biological Association of the United Kingdom,102(8), 613–626. 10.1017/S0025315422001011

Sibley, E. C. P., Madgett, A. S., Elsdon, T. S., Marnane, M. J., Harvey, E. S., Songploy, S., Kettradad, J., and Fernandes, P. G. (2023). An acoustic-optic comparison of fish assemblages at a Rigs-to-Reefs habitat and coral reef in the Gulf of Thailand. Estuarine, Coastal and Shelf Science, 295. 10.1016/j.ecss.2023.108552

Simon, T., Joyeux, J. C., and Pinheiro, H. T. (2013). Fish assemblages on shipwrecks and natural rocky reefs strongly differ in trophic structure. Marine Environmental Research, 90, 55–65. 10.1016/j.marenvres.2013.05.012

Streich, M. K., Ajemian, M. J., Wetz, J. J., and Stunz, G. W. (2017). A comparison of fish community structure at mesophotic artificial reefs and natural banks in the western gulf of mexico. Marine and Coastal Fisheries, 9(1), 170–189. 10.1080/19425120.2017.1282897

Streit, R. P., Hoey, A. S., and Bellwood, D. R. (2015). Feeding characteristics reveal functional distinctions among browsing herbivorous fishes on coral reefs. Coral Reefs, 34(4), 1037–1047. 10.1007/S00338-015-1322-Y/METRICS

Vehtari, A., Gelman, A., and Gabry, J. (2017). Practical Bayesian model evaluation using leave-one-out cross-validation and WAIC. Statistics and Computing, 27, 1413–1432. 10.1007/s11222-016-9696-4

Yeemin, T., Sutthacheep, M., and Pettongma, R. (2006). Coral reef restoration projects in Thailand. Ocean and Coastal Management, 49(9–10), 562–575. 10.1016/J.OCECOAMAN.2006.06.002

White, M., Bashmachnikov, I., Arístegui, J., and Martins, A. (2007). Physical processes and seamount productivity. In T. J. Pitcher, T. Morato, P. J. B. Hart, M. R. Clark, N. Haggan, and R. S. Santos (Eds.), Seamounts: Ecology, Fisheries and Conservation (pp. 62–84). Blackwell Publishing

Wickham, H., Averick, M., Bryan, J., Chang, W., McGowan, L. D., François, R., Grolemund, G., Hayes, A., Henry, L., Hester, J., Kuhn, M., Lin Pedersen, T., Miller, E., Bache, S. M., Müller, K., Ooms, J., Robinson, D., Seidel, D. P., Spinu, V., … Yutani, H. (2019). Welcome to the tidyverse. Journal of Open Source Software, 4(43), 1686. 10.21105/joss.01686

