## Supplemental Tables S1-S3 for "Shipwrecks can mirror predator assemblages of pelagic pinnacles but may lack the trophic balance of natural reef environments"

Table S1  List of reef-associated taxa recorded during surveys, including common names, genus and species-level identifications (where possible), and assigned functional group classifications. Species chosen are the most commonly spotted in the waters surrounding Koh Tao, span a range of trophic levels and ecological roles, and include both site-attached and mobile predators.

| **Common Name** | **Genus** | **Species** | **Functional Group** |
| --- | --- | --- | --- |
| Parrotfish | *Scarus* | spp. | Grazer |
| Rabbitfish | *Siganus* | spp. | Grazer |
| Butterflyfish | *Chaetodon* | spp. | Grazer |
| Angelfish | *Pomacanthus* | spp. | Invertivore |
| Cleaner Wrasse | *Labroides* | *dimidiatus* | Invertivore |
| Batfish | Ephippidae | spp. | Invertivore |
| Thicklip Wrasse | *Hemigymnus* | *melapterus* | Invertivore |
| Red Breast Wrasse | *Cheilinus* | *fasciatus* | Invertivore |
| Slingjaw Wrasse | *Epibulus* | *insidiator* | Invertivore |
| Sweetlips | *Diagramma / Plectorhinchus* | spp. | Invertivore |
| Squirrelfish and Soldierfish | Holocentridae | spp. | Invertivore |
| Triggerfish | Balistidae | spp. | Invertivore |
| Porcupinefish and Pufferfish | *Diodon / Tetraodon* | spp | Invertivore |
| Ray | *Taeniura / Neotrygon* | spp. | Mesopredator |
| Brown Stripe Snapper | *Lutjanus* | *vitta* | Mesopredator |
| Russell's Snapper | *Lutjanus* | *russellii* | Mesopredator |
| Large (>30cm) Snapper | *Lutjanus* | spp. | HTLP |
| Eel | *Gymnothorax* | spp. | Mesopredator |
| Trevally | *Caranx* | spp. | HTLP |
| Emperorfish | *Lethrinus* | spp. | Mesopredator |
| Small (<30cm) Grouper | *Cephalopholis / Epinephelus* | spp. | Mesopredator |
| Large (>30cm) Grouper | *Epinephelus* | spp. | HTLP |
| Barracuda | *Sphyraena* | spp. | HTLP |

Table S2 Predicted mean abundances (with 95% credible intervals) of 18 reef fish taxa across three reef classifications: shipwreck, fringing reef, and pelagic pinnacle. Estimates are derived from a Bayesian multivariate negative binomial model with reef classification as a fixed effect and site as a random intercept. Taxa are grouped by functional role (grazer, invertivore, mesopredator, HTLP), and values reflect the expected total count per survey.

| **Taxa** | **Functional Group** | **Shipwreck** | **Fringing** | **Pinnacle** |
| --- | --- | --- | --- | --- |
| *Scarus* spp. | Grazer | 7.47 (6.39–8.78) | 48.42 (43.69–53.81) | 19.35 (17.62–21.34) |
| *Siganus* spp. | Grazer | 40.88 (34.36–49.05) | 38.19 (33.59–43.67) | 59.39 (53.02–66.87) |
| *Chaetodon* spp. | Grazer | 11.21 (9.80–12.88) | 44.57 (40.79–48.84) | 31.84 (29.37–34.62) |
| *Pomacanthus* spp. | Invertivore | 1.01 (0.80–1.26) | 2.57 (2.28–2.90) | 3.24 (2.92–3.59) |
| *Labroides dimidiatus* | Invertivore | 0.90 (0.68–1.17) | 8.86 (7.77–10.12) | 8.70 (7.74–9.81) |
| Ephippidae spp. | Invertivore | 0.20 (0.10–0.41) | 0.34 (0.22–0.54) | 1.87 (1.35–2.66) |
| *Hemigymnus melapterus* | Invertivore | 0.07 (0.03–0.14) | 6.13 (5.31–7.09) | 2.13 (1.84–2.48) |
| *Cheilinus fasciatus* | Invertivore | 2.44 (1.92–3.12) | 12.92 (11.12–15.12) | 6.43 (5.60–7.45) |
| *Epibulus insidiator* | Invertivore | 0.16 (0.09–0.26) | 6.01 (5.19–7.01) | 1.01 (0.84–1.21) |
| *Diagramma / Plectorhinchus* spp. | Invertivore | 29.36 (21.39–40.85) | 7.35 (5.84–9.42) | 15.23 (12.46–18.93) |
| Holocentridae spp. | Invertivore | 8.21 (5.56–12.77) | 8.29 (6.27–11.17) | 23.95 (18.68–31.17) |
| Balistidae spp. | Invertivore | 0.64 (0.48–0.84) | 2.49 (2.18–2.84) | 1.09 (0.94–1.26) |
| *Lutjanus* spp. (<30cm) | Mesopredator | 105.95 (74.84–156.53) | 29.33 (22.74–38.45) | 100.88 (80.22–128.99) |
| *Cephalopolis / Epinephelus* spp. (<30cm) | Mesopredator | 4.46 (3.77–5.30) | 33.18 (29.97–36.79) | 36.20 (33.10–39.74) |
| *Lethrinus* spp. | Mesopredator | 5.35 (4.17–7.00) | 6.06 (5.07–7.32) | 13.00 (11.14–15.33) |
| *Lutjanus* spp. (>30cm) | HTLP | 12.82 (9.58–17.92) | 0.74 (0.57–0.99) | 4.47 (3.66–5.55) |
| *Epinephelus* spp. (>30cm) | HTLP | 6.11 (5.06–7.48) | 5.69 (4.95–6.57) | 8.89 (7.88–10.10) |
| *Caranx* spp. | HTLP | 8.02 (6.28–10.42) | 4.67 (3.89–5.67) | 18.81 (16.10–22.19) |

Table S3 Posterior probabilities of pairwise differences in predicted log-abundance for 18 reef taxa across reef classifications (shipwreck, fringing reef, pinnacle). Values are derived from a Bayesian multivariate negative binomial model. The table reports the posterior median, 95% credible interval (CI), and the proportion of posterior samples supporting positive or negative differences.

| **Taxa** | **Functional Group** | **Comparison** | **Median** | **CI_lower_** | **CI_upper_** | **P(β>0)** | **P(β<0)** |
| --- | --- | --- | --- | --- | --- | --- | --- |
| *Scarus spp.* | Grazer | Fringing – Shipwreck | **1.87** | 1.68 | 2.06 | 1 | 0 |
|  |  | Pinnacle – Fringing | **-0.917** | -1.06 | -0.78 | 0 | 1 |
|  |  | Pinnacle – Shipwreck | **0.953** | 0.77 | 1.14 | 1 | 0 |
| *Siganus spp.* | Grazer | Fringing – Shipwreck | **-0.068** | -0.287 | 0.153 | 0.272 | 0.728 |
|  |  | Pinnacle – Fringing | **0.441** | 0.27 | 0.612 | 1 | 0 |
|  |  | Pinnacle – Shipwreck | **0.373** | 0.152 | 0.579 | 0.999 | 0.001 |
| *Chaetodon spp.* | Grazer | Fringing – Shipwreck | **1.38** | 1.22 | 1.54 | 1 | 0 |
|  |  | Pinnacle – Fringing | **-0.335** | -0.462 | -0.213 | 0 | 1 |
|  |  | Pinnacle – Shipwreck | **1.04** | 0.887 | 1.2 | 1 | 0 |
| *Pomacanthus spp.* | Invertivore | Fringing – Shipwreck | **0.936** | 0.677 | 1.2 | 1 | 0 |
|  |  | Pinnacle – Fringing | **0.231** | 0.074 | 0.387 | 0.999 | 0.002 |
|  |  | Pinnacle – Shipwreck | **1.16** | 0.92 | 1.42 | 1 | 0 |
| *Labroides dimidiatus* | Invertivore | Fringing – Shipwreck | **2.29** | 1.99 | 2.59 | 1 | 0 |
|  |  | Pinnacle – Fringing | **-0.017** | -0.194 | 0.162 | 0.426 | 0.574 |
|  |  | Pinnacle – Shipwreck | **2.27** | 1.98 | 2.57 | 1 | 0 |
| *Ephippidae spp.* | Invertivore | Fringing – Shipwreck | **0.539** | -0.281 | 1.35 | 0.908 | 0.092 |
|  |  | Pinnacle – Fringing | **1.7** | 1.14 | 2.25 | 1 | 0 |
|  |  | Pinnacle – Shipwreck | **2.24** | 1.46 | 2.99 | 1 | 0 |
| *Hemigymnus melapterus* | Invertivore | Fringing – Shipwreck | **4.52** | 3.81 | 5.4 | 1 | 0 |
|  |  | Pinnacle – Fringing | **-1.05** | -1.26 | -0.846 | 0 | 1 |
|  |  | Pinnacle – Shipwreck | **3.47** | 2.75 | 4.36 | 1 | 0 |
| *Cheilinus fasciatus* | Invertivore | Fringing – Shipwreck | **1.67** | 1.37 | 1.97 | 1 | 0 |
|  |  | Pinnacle – Fringing | **-0.698** | -0.908 | -0.484 | 0 | 1 |
|  |  | Pinnacle – Shipwreck | **0.973** | 0.672 | 1.26 | 1 | 0 |
| *Epibulus insidiator* | Invertivore | Fringing – Shipwreck | **3.65** | 3.12 | 4.25 | 1 | 0 |
|  |  | Pinnacle – Fringing | **-1.79** | -2.03 | -1.55 | 0 | 1 |
|  |  | Pinnacle – Shipwreck | **1.86** | 1.32 | 2.46 | 1 | 0 |
| *Diagramma/ Plectorhinchus spp.* | Invertivore | Fringing – Shipwreck | **-1.38** | -1.79 | -0.997 | 0 | 1 |
|  |  | Pinnacle – Fringing | **0.731** | 0.423 | 1.03 | 1 | 0 |
|  |  | Pinnacle – Shipwreck | **-0.651** | -1.04 | -0.284 | 0 | 1 |
| *Holocentridae spp.* | Invertivore | Fringing – Shipwreck | **0.009** | -0.503 | 0.499 | 0.515 | 0.485 |
|  |  | Pinnacle – Fringing | **1.06** | 0.666 | 1.45 | 1 | 0 |
|  |  | Pinnacle – Shipwreck | **1.08** | 0.574 | 1.54 | 1 | 0 |
| *Balistidae spp.* | Invertivore | Fringing – Shipwreck | **1.36** | 1.06 | 1.67 | 1 | 0 |
|  |  | Pinnacle – Fringing | **-0.825** | -1.02 | -0.628 | 0 | 1 |
|  |  | Pinnacle – Shipwreck | **0.536** | 0.225 | 0.857 | 1 | 0 |
| *Lutjanus (<30cm) spp.* | Mesopredator | Fringing – Shipwreck | **-1.29** | -1.76 | -0.851 | 0 | 1 |
|  |  | Pinnacle – Fringing | **1.24** | 0.888 | 1.58 | 1 | 0 |
|  |  | Pinnacle – Shipwreck | **-0.051** | -0.508 | 0.374 | 0.405 | 0.595 |
| *Lethrinus spp.* | Mesopredator | Fringing – Shipwreck | **0.121** | -0.204 | 0.442 | 0.768 | 0.232 |
|  |  | Pinnacle – Fringing | **0.767** | 0.52 | 1 | 1 | 0 |
|  |  | Pinnacle – Shipwreck | **0.889** | 0.581 | 1.19 | 1 | 0 |
| *Cephalopholis/ Epinephelus spp.* | Mesopredator | Fringing – Shipwreck | **2.01** | 1.81 | 2.2 | 1 | 0 |
|  |  | Pinnacle – Fringing | **0.088** | -0.05 | 0.226 | 0.89 | 0.11 |
|  |  | Pinnacle – Shipwreck | **2.09** | 1.9 | 2.28 | 1 | 0 |
| *Lutjanus (>30cm) spp.* | HTLP | Fringing – Shipwreck | **-2.85** | -3.28 | -2.44 | 0 | 1 |
|  |  | Pinnacle – Fringing | **1.8** | 1.46 | 2.12 | 1 | 0 |
|  |  | Pinnacle – Shipwreck | **-1.06** | -1.44 | -0.692 | 0 | 1 |
| *Caranx spp.* | HTLP | Fringing – Shipwreck | **-0.542** | -0.867 | -0.229 | 0.001 | 0.999 |
|  |  | Pinnacle – Fringing | **1.39** | 1.15 | 1.64 | 1 | 0 |
|  |  | Pinnacle – Shipwreck | **0.849** | 0.544 | 1.15 | 1 | 0 |
| *Epinephelus (>30cm)/ Plectropomus spp.* | HTLP | Fringing – Shipwreck | **-0.074** | -0.318 | 0.161 | 0.27 | 0.73 |
|  |  | Pinnacle – Fringing | **0.446** | 0.26 | 0.634 | 1 | 0 |
|  |  | Pinnacle – Shipwreck | **0.371** | 0.138 | 0.599 | 0.999 | 0.001 |
